# Learning copy number dependent variation in single tumour cell transcriptomes with deep generative models

**DOI:** 10.1101/2025.04.21.649842

**Authors:** Alina Selega, Hassaan Maan, Chengxin Yu, Tiak Ju Tan, Kieran R Campbell

## Abstract

Large-scale copy number aberrations are a hallmark of many cancers, showing widespread associations with tumour cell phenotypes, therapy resistance, and patient outcomes. Several existing methods can infer copy number profiles from single-cell RNA-sequencing data. However, there is currently no methodology to separate the effects of the copy number state on single tumour cell phenotypes from the copy number-independent variation. We present ISOMERIC, an unsupervised deep learning framework to learn disentangled representations of single-cell expression data that are dependent on or independent of copy number profiles. We apply ISOMERIC to multiple cancer types and examine how copy number shapes transcriptional states. We find distinct associations between copy number-dependent and independent variation and clinical subtypes in multiple cancer types and link such variation to tumour microenvironment phenotypes. Together, this establishes a principled framework for untangling the effects of genetic and intrinsic variation on tumour transcriptomes and the consequences on clinically meaningful phenotypes.

## Introduction

Somatic copy number aberrations are one of the hallmarks of cancer^1^, and many tumours are typically composed of genetically heterogeneous cellular populations sharing distinct copy number profiles—also known as clones^2^. Clones often have different proliferative and treatment resistant capabilities and their presence, absence, or interaction may dramatically alter the course of the disease and patient outcomes^3–6^. Thus, mapping associations between a tumour cell’s genetic aberrations and its gene expression phenotype may help identify clonal populations associated with poor clinical outcomes and uncover the aberrant mechanisms that confer malignant properties to cancer cells.

Currently, only a handful of technologies exist that simultaneously assay RNA and DNA in the same single cells to simultaneously quantify a cell’s copy number and transcriptional state^7–9^. However, these technologies have not yet achieved the high throughput and scalability of single-cell RNA-sequencing (scRNA-seq), an assay profiling whole-transcriptome expression in single cells. Due to this scalability and the ability to generate a cell-level readout of heterogeneity in a sample, an extensive compendium of scRNA-seq data from cancer studies are publicly available to the community. Several computational methods have been developed to integrate scRNA-seq and copy number data^10,11^, infer copy number calls from scRNA-seq^12–14^, or jointly analyze the two modalities^15^. Together, these technologies have confirmed that cellular genomic copy number is a crucial factor affecting the expression profile of individual tumour cells, which often cluster based on the patterns of their copy number aberrations^16^.

However, this dependency confounds any analysis seeking to identify shared or distinct transcriptional states either driven by or independent of cellular genomic copy number and subsequently relate them to clinically-relevant measures. Several methods have been introduced to isolate salient dimensions of variation from single-cell and spatial data, either under the assumption of specific perturbations^17^ or co-measured multi-modal data^18^. However, no existing method integrates paired scRNA-seq and copy number profiles while explicitly separating copy number-dependent and -independent variation in tumour cell phenotypes. To address this, we present ISOMERIC (d**IS**entangling c**O**py nu**M**ber d**E**pendent va**R**iation **I**n trans**C**riptomes), an adversarial variational autoencoder that decomposes expression profiles of single tumour cells into copy number dependent and independent components. We trained ISOMERIC on the expression from 189,027 single tumour cells from a total of 10 publicly available cohorts from pancreatic, breast, ovarian, and lung cancers. We found multiple copy number-dependent latent dimensions that encoded known mutational signatures and transcriptional subtypes in ovarian and pancreatic cancers. We uncovered multiple copy number dependent and independent sources of variation that are coupled to tumour microenvironment composition across all four cancers we considered. Overall, we demonstrate that ISOMERIC learns latent representations that recapitulate clinically-relevant signatures and show how they can be used to identify associated loci and gene programs.

## Results

### ISOMERIC decomposes single-cell expression into copy number associated and independent components

To separate copy number dependent and independent variation in scRNA-seq of tumour cells, we adapted existing variational autoencoder (VAE) models^19^. VAEs learn a lower-dimensional representation termed a *latent space* by compressing the input expression data via an encoder network, then decompressing via a decoder network to reconstruct the input. Such models are trained by minimizing a loss that probabilistically measures the discrepancy between the input and reconstruction. VAEs and their variants have been extensively applied to both scRNA-seq and multi-modal datasets^17,18,20,21^.

Inspired by approaches for disentangled representation learning^22^, we adapted the VAE model to simultaneously learn two latent spaces: one that represents variation in scRNA-seq data associated with copy number that we term *CNA*_*d*_ (CNA-dependent), and one that represents variation independent of copy number that we term *CNA*_*i*_ (CNA-independent) (**Fig. 1A** and **Methods**). ISOMERIC achieves this decomposition by incorporating two additional components into an scRNA-seq autoencoder: a CNA decoder network that learns to predict copy number from CNA_d_ and a second neural network that learns to predict copy number from CNA_i_ that is adversarially trained. As input, ISOMERIC requires both scRNA-seq from tumour cells and paired cell-by-genomic region copy number, either inferred from scRNA-seq data with methods such as inferCNV^12^ or measured jointly using assays such as G&T-seq.

**Figure 1.**
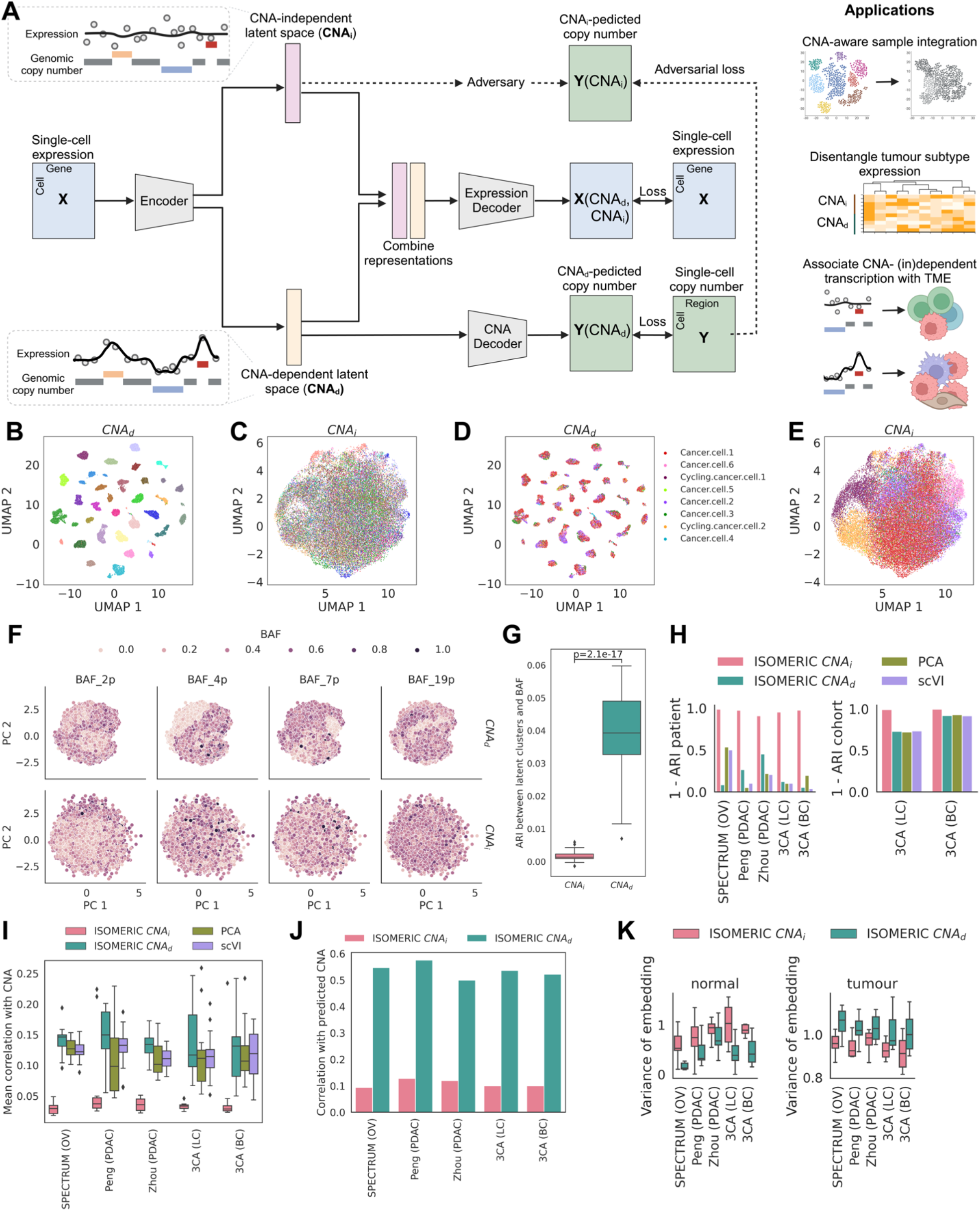
Learning copy number dependent variation from single-cell transcriptomes. **A** The ISOMERIC model architecture (left) and model applications (right). **B-E** UMAP projections of the SPECTRUM dataset using latent spaces coloured by patient identity for CNA_d_ (**B**) and CNA_i_ (**C**), and coloured by original publication’s cancer cell cluster for CNA_d_ (**D**) and CNA_i_ (**E**). **F** PCA projections of SPECTRUM dataset on CNA_d_ and CNA_i_ latent spaces colored by BAF score on selected chromosome arms. **G** Adjusted Rand Index (ARI) between CNA_d_ and CNA_i_ clusters and discretized BAF scores across all chromosome arms. Statistical test performed using two-sided t-test **H** Mixing of latent spaces learned by ISOMERIC and two baselines (scVI) for both patient ID (left) and cohort ID (right). **I** Correlation between tumour cells latent representation and cellular copy number. **J** Correlation between true and reconstructed copy number profiles. **K** Variance of embeddings in ISOMERIC CNA_d_ and CNA_i_ for normal cells and tumour cells.

We developed a benchmarking pipeline to validate the utility of our model using publicly available scRNA-seq datasets from cancer-focused studies that included multi-patient cohorts and featured copy number analysis. To do so we collated two pancreatic ductal adenocarcinoma (PDAC) cohorts: Zhou^23^ and Peng^24^, one ovarian cancer (OV) cohort called SPECTRUM^25^, four lung cancer (LC) cohorts^26–29^ in the 3CA repository^30^ and three breast cancer (BC) cohorts^27,31,32^ also in the 3CA repository. Each dataset was filtered to contain malignant cells based on observed copy number variation (**Methods**).

With paired CNAs inferred from each dataset, we trained ISOMERIC on the five datasets separately for each cancer type (**Methods**). We first created Uniform Manifold Approximation and Projection (UMAP) plots^33^ of the tumour cell embeddings in each of the latent spaces, CNA_i_ and CNA_d_. Upon visual inspection of the UMAP plots projecting the SPECTRUM (OV) data, we found that the CNA_d_ space appeared separated by patient ID (**Fig. 1B**) while the CNA_i_ space appeared well-mixed (**Fig. 1C**). This matches the intuition that patients will have distinct copy number profiles resulting in unique expression programs captured by CNA_d_, while copy number independent transcription captured by CNA_i_ should be recurrent between patients. Consistent with this, CNA_i_ largely captured variation associated with cell cycle as made apparent by overlaying the original cell cluster annotations that demonstrate visual separation of two clusters of cycling cancer cells in CNA_i_ but not CNA_d_ (**Fig. 1D-E**). We additionally used the cell-specific B-allele frequency (BAF) scores from the SPECTRUM (OV) metadata to investigate how this orthogonal measure of single-cell genetic variation was represented in the latent spaces. We found CNA_d_ to be visually separated by BAF scores in contrast to CNA_i_ which was more mixed on a principal components analysis (PCA) projection of the data (**Fig. 1F**). We explicitly quantified this by clustering each latent space and measuring the agreement between the clusters and quantized BAF values. Indeed, CNA_d_ was significantly more associated with BAF (p<0.01, **Fig. 1G**).

We next contrasted how existing methods for scRNA-seq dimensionality reduction encode copy number information within their latent spaces compared to ISOMERIC. We implemented two common baselines: scVI^21^, a commonly used VAE-based method for scRNA-seq data, and PCA. We trained both benchmarks on the tumour cell scRNA-seq data from the five considered datasets (**Methods**). We then investigated several metrics for all three methods: (i) concordance between latent space clusters and patient identity, (ii) concordance between latent space clusters and cohort identity (for pooled breast and lung cancer datasets), and (iii) the correlation between latent dimensions and the input copy number profiles (**Methods**). As expected, ISOMERIC CNA_i_ was strongly independent of patient identity, despite not explicitly having patient specific information, while ISOMERIC CNA_d_ along with baselines showed larger though variable dependence (**Fig. 1H**, left). Similarly, ISOMERIC CNA_d_ was less dependent on dataset than ISOMERIC CNA_i_ and baselines (**Fig. 1H**, right), implying the ability to learn recurrent copy number independent transcriptional states across cohorts. In addition, as expected ISOMERIC CNA_d_ had on average the highest correlation with single-cell copy number, followed by the two benchmarks, and then ISOMERIC CNA_i_ with the lowest correlation (**Fig. 1I**). The capacity of CNA_d_ and CNA_i_ to represent copy number was further corroborated by high/low correlation between true and predicted CNA as decoded from each latent space **(Fig. 1J)**. Together, this implies latent spaces from existing methods learn a mix of copy number dependent and independent transcriptional variation, which can be effectively decomposed using ISOMERIC. Finally, we embedded normal epithelial cells from the same patients into both latent spaces and measured the variance of those embeddings (**Methods**), with the intuition that normal cells should occupy a similar part of CNA_d_ due to no aneuploidy but still vary in a copy number independent manner in CNA_i_. As anticipated, normal cells had lower variance of embeddings in CNA_d_ than CNA_i_ across all datasets (**Fig. 1K**, left), which is notable given ISOMERIC was not trained on any non-malignant cells. Interestingly, tumour cells had more similar variance across latent spaces, with on average lower variance in CNA_i_, highlighting more CNA-dependent variation in their expression (**Fig. 1K**, right). Taken together, these results show that ISOMERIC decomposes expression profiles of single tumour cells into components that are associated with and independent of their copy number.

### ISOMERIC latent spaces encode cancer subtypes

We next evaluated whether the ISOMERIC latent spaces were associated with clinical variables and states. We leveraged the SPECTRUM (OV) dataset of single-cell tumour profiling of 42 high grade serous ovarian tumours with rich metadata including tumour site and mutational signature annotations. We computed sample-level mean embeddings in each latent dimension of CNA_i_ and CNA_d_. We found multiple significant associations with the sample tumour site and genomically defined patient homologous recombination deficient (HRD) and CCNE1 amplification-associated foldback inversion (FBI) clinical subtypes. Specifically, we found two CNA_i_ dimensions significantly associated with the tumour site (p_adj_<0.05 with Benjamini-Hochberg correction) but no CNA_d_ dimensions associated with site (**Fig. 2A**). Conversely, we found five CNA_d_ dimensions and one CNA_i_ dimension significantly associated (p_adj_<0.05) with the HRD/FBI signature (**Fig. 2B**). Examining the boxplots of sample-level embeddings for each HRD/FBI signature revealed that the four CNA_d_ dimensions (w7, w9, w6, w1) appear to separate two HRD subtypes, BRCA1-associated tandem duplications (HRD-Dup) and BRCA2-associated interstitial deletions (HRD-Del). On the other hand, the CNA_d_ dimension with the strongest association (w5) more strongly separated the FBI and HRD signatures (**Fig. 2C**). We then examined which chromosome bands correlated with the strongest signature-associated CNA_d_ dimension (w5). We found high correlations between the embeddings and the copy number in many genomic regions, most notably within chromosomes 2 and 12, with the maximum association at chromosome band 12q13.12 (**Fig. 2D**), a well known site of tumour suppressor genes in ovarian cancer^34^. Together, these findings suggest tumour evolution minimally shapes transcriptomes in a tumour site-dependent manner in ovarian cancer but significantly shapes the transcriptome dependent on clinically-actionable subtypes.

**Figure 2.**
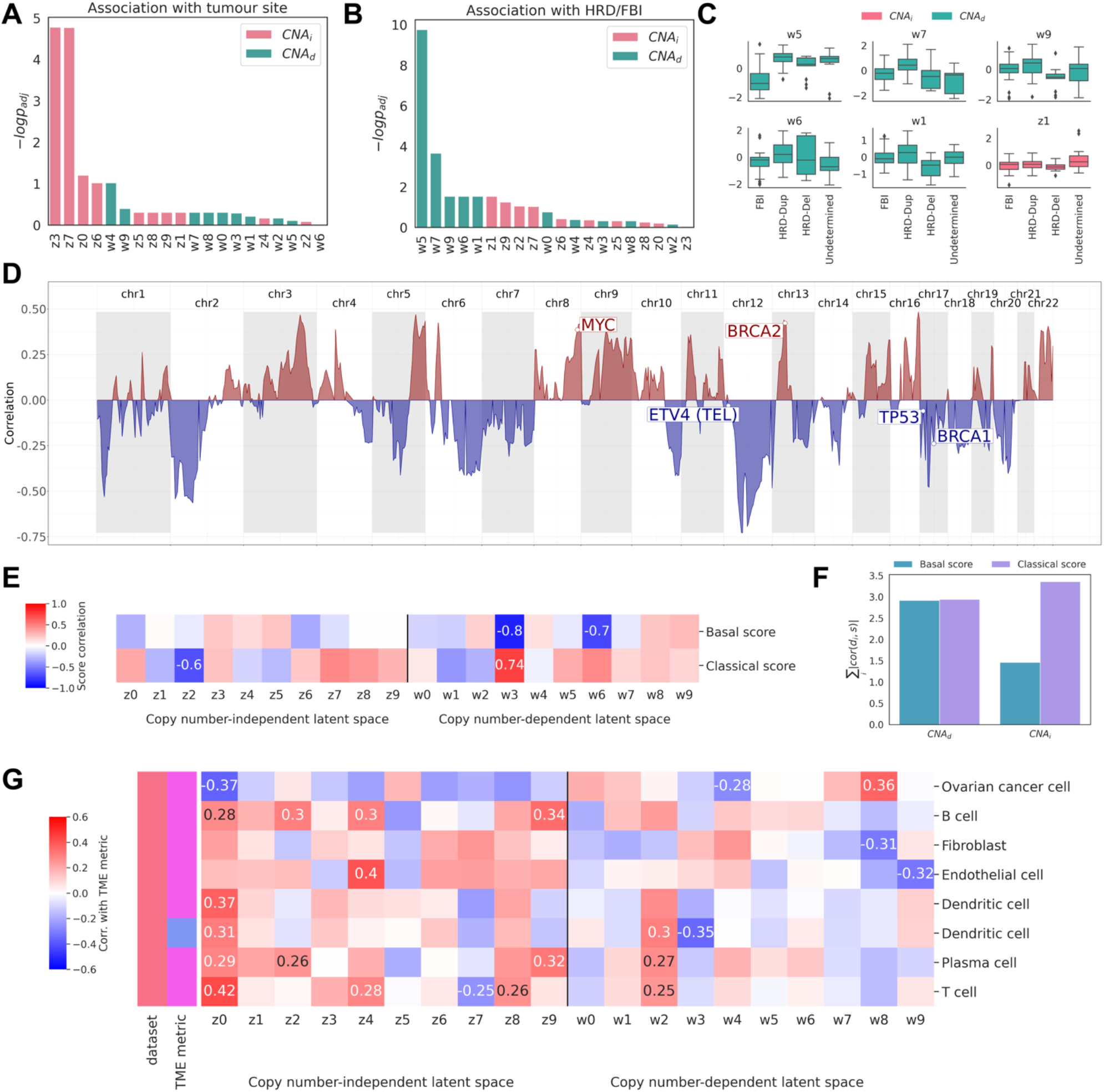
ISOMERIC learns latent spaces associated with tumour subtypes and TME composition. **A** Association between sample-mean cell embeddings in each ISOMERIC latent dimension and tumour site for the SPECTRUM (OV) dataset. P-values were multiple test corrected. **B** As **A** for ovarian mutational signature subtype. **C** Mean ISOMERIC cell embeddings dependent on mutational signature subtype for six select dimensions. **D** Correlation between the chromosome band copy number and one latent dimension of the CNA_d_ space (w5). **E** Correlation between patient-level embeddings of PDAC cells in each ISOMERIC dimension and patient PDAC subtype scores. Significantly associated dimensions (p_adj_<0.05) are annotated. **F** The sum of absolute correlations across dimensions in each ISOMERIC latent space for each subtype. **G** Correlation between cell type proportions (TME metrics) and mean ISOMERIC tumour cell embeddings in each latent dimension for SPECTRUM (OV). Significant correlations (p_adj_ < 0.05) are annotated.

We next leveraged ISOMERIC to quantify copy number dependence on single-cell variation of two expression subtypes in PDAC. PDAC tumours are frequently classified into *classical* and *basal* subtypes based on bulk RNA-seq, with classical patients exhibiting improved survival compared to basal^35^. Using a recent single-cell atlas Zhou^23^ encompassing 31 patients and 232,764 cells we had previously annotated^36^, we used the published gene sets associated with the basal and classical subtypes to compute a subtype score for each tumour cell based on its expression (**Methods**). Correlating these with ISOMERIC latent dimensions, we found two CNA_d_ and one CNA_i_ dimensions were significantly associated (p_adj_<0.05) with the basal or classical score, with a specific CNA_d_ dimension representing subtypes as a spectrum (CNA_d_high strongly correlated with the classical subtype and CNA_d_low with the basal subtype, **Fig. 2E**). When considering strength of association across dimensions by summing the absolute values of the correlation coefficients, all CNA_d_ dimensions were associated with both subtypes, while CNA_i_ dimensions more strongly correlated with the classical score (**Fig. 2F**). Together, our analysis suggests that copy number variation may significantly drive single-cell transcriptional variation leading to the poor outcome basal PDAC subtype, while both copy number dependent and independent variation jointly contributes to the classical subtype.

### ISOMERIC associates latent tumour cell states with tumour microenvironment composition

We next investigated whether ISOMERIC latent spaces can be used to associate copy number dependent and independent variation with tumour microenvironment (TME) composition. We considered two metrics characterizing the TME: (i) the proportion a given cell type makes up of all cells in an individual sample and (ii) the proportion a given immune cell type makes up of all immune cells in a sample (**Methods**). We computed these metrics for all cell types present in each dataset and correlated these metrics with the average latent embeddings in each dimension to examine if any dimensions encode the TME makeup (**Methods**).

We found multiple cell types whose overall cell type proportion was significantly associated (p_adj_ < 0.05, Benjamini-Hochberg corrected) with ISOMERIC latent dimensions for all datasets (**S. Fig. 1**), except for 3CA (BC) which was only associated with an epithelial cell type. Notably, the abundance of different TME cell types was differentially correlated with either copy number dependent or independent tumour cell variation in the ovarian dataset. B cell proportions within the TME were uniquely correlated with multiple copy number independent tumour cell latent dimensions, fibroblast proportions were uniquely correlated with a copy number dependent dimension, while other cell type proportions were determined by tumour transcriptomic variation that was both copy number dependent and intrinsic (**Fig. 2G**). Together, these analyses highlight the ability to link tumour microenvironment composition to tumour cell phenotypes and lay a foundation for further cell-cell communication analysis^37^ and future functional studies.

## Discussion

In this work we introduced ISOMERIC, an adversarial deep learning framework that learns disentangled representations of tumour cell transcriptomes, separating tumour cell phenotypes into components that are associated with or independent of the copy number. We showed that ISOMERIC effectively disentangles copy number variation from the transcriptome and in doing so learns representations that capture clinically-relevant variation, both related to molecular subtypes and TME composition. Notably, ISOMERIC achieves batch integration in the CNA-independent latent space as a by-product of the training procedure ensuring copy number independence. We further introduced a set of analyses for the interpretation of individual latent dimensions that may help identify genomic loci or gene programs of interest when analyzing tumour and tumour-microenvironment interactions.

However, multiple challenges remain and may act as future research directions. Copy number aberrations appear at different length scales, and while we engineered features at the cytoband-scale, this may miss focal amplifications and deletions or place too much weight on large scale copy number aberrations in the latent space. Likewise, multiple factors come together to influence tumour cell expression beyond genomic copy number, including epigenetics, exogenous perturbations, and cellular environments of the tumour microenvironment. As paired data jointly measuring these becomes available, future methods may try to fully explain observed variation as a function of all possible causes. Finally, genomic copy number influences expression through both *in-cis* and *in-trans* action across the genome^38^. While our proposed framework does not explicitly capture this, future methods may actively try to explain latent representations in terms of such mechanisms, providing insights to gene networks driving malignant states and ultimately possible therapies.

## Methods

### Model

ISOMERIC embeds single tumour cells into two latent spaces, one of which represents variation in both cell’s expression and CNA profile (CNA_d_, with latent dimensions denoted w_j_) and the other, which represents variation in expression only (CNA_i_, with latent dimensions denoted z_j_). CNA_d_ stands for copy number-dependent, CNA_i_ – for copy number-independent, and j ∈ 1, …, *D*, where *D* is the dimensionality of each latent space.

We denote the scRNA-seq data matrix as **X**, which has *N* rows corresponding to cells and *G* columns corresponding to genes. The second input matrix **Y** contains single-cell CNA profiles for the same cells and has *R* columns corresponding to genomic regions. These CNA profiles can be summarized to varying region sizes, such as chromosome arms (with *R*=44, excluding chromosomes X, Y) or Giesma chromosome band-level aggregated CNA profiles (with *R*=811 Giesma bands covering 1-22 chromosomes). See *Hyperparameter tuning* and *Copy number profile generation* for further details.

We decompose the joint distribution over the input data (**x**,**y**) and latent variables (**z, w**) as:

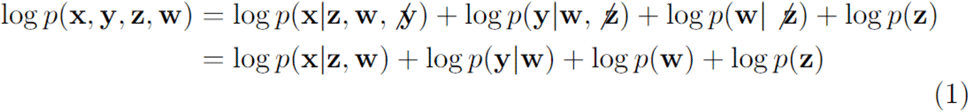

This assumes the conditional independence relationships *x* ⊥ *y* | *z*,*w*,*y* ⊥ *z*,*w* ⊥ *z* In other words, given a cell’s latent representations, its CNA profile does not add any new information to its expression, CNA-independent latent space **z** is independent of CNA profiles **y**, and the two latent spaces **w** and **z** are independent of each other.

Note that this decomposition is different from Klys et al.^22^ where they condition the prior over **w** on **y** and model different **w** subspaces for each category of **y**. We adapted our model for a continuous **y (**indicating per-cell CNA profiles) and instead modelled the dependency between **w** and **y** using a CNA decoder, which learns to reconstruct **y** from a posterior sample **w** (see below). This objective helps us ensure that the CNA_d_ latent space learns to represent salient information about tumour cell CNA profiles.

We train our model using the variational lower bound on the marginal log likelihood of the data and imposing independence between **z** and **y**. Given an approximate joint posterior *q*(**z**,**w**|**x**,**y**) that we aim to learn, we can derive the evidence lower bound (ELBO) traditionally used for VAEs:

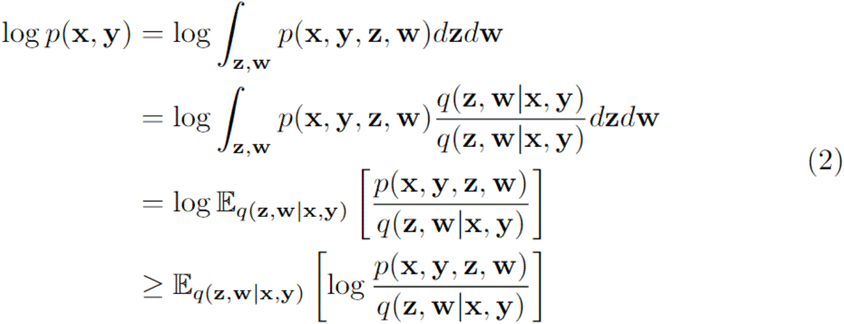

Using our joint distribution decomposition and taking a negative of the above gives an upper bound on *p*(**x**,**y**) that we seek to minimize:

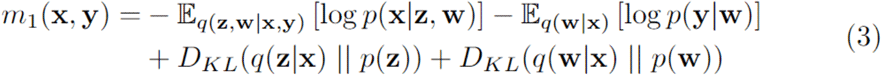

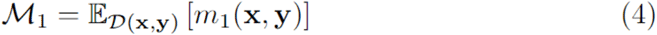

Our expression for the ELBO contains a reconstruction error term for expression **x** from both latent spaces **z** and **w**; a KL divergence term for the posterior over each of the latent variables; and a reconstruction error term for CNA profiles **y** when decoded from the CNA_d_ latent space (**w**). The priors over **w** and **z** are modelled as standard normal. To ensure that **y** is independent of **z**, we add the following term to the objective to discourage the ability to predict CNA profiles **y** from the CNA_i_ latent space (**z**):

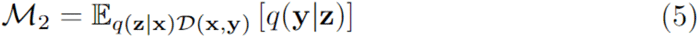

Note that *adding M*_*2*_ to our minimized objective amounts to *minimizing* the likelihood of reconstructed **y** under *q*(**y**|**z**), achieving our goal of making it hard to predict CNA profiles from **z**. As in Klys et al.^22^, *M*_*2*_ represents the adversarial component of the network, where the adversary *q*(**y**|**z**) tries to predict CNA profiles **y** from a posterior sample **z**, while the encoder *q*(**z**|**x**) tries to generate **z** to prevent this. Our final objective consists of two parts, which we train jointly, as done in Klys et al. *β*_*i*_ for i=1,2,3 weights are treated as hyperparameters.

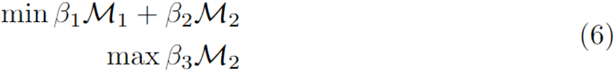

We model the likelihood functions for the expression decoder *p*(**x**|**z**,**w**), the CNA decoder *p*(**y**|**w**) and the adversary *q*(**y**|**z**) as Gaussian distributions with constant variance of 1 for each feature (gene, chromosome arm, or chromosome band; see below for more details). We have also considered a negative binomial likelihood function for the expression data but found that Gaussian likelihood performed better (see *Hyperparameter Tuning*). The means of the distributions are estimated with corresponding neural networks.

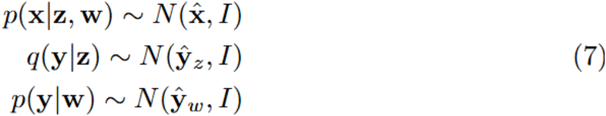

When modelling datasets that were pooled over multiple cohorts, we introduce an indicator variable **c** to denote the cohort. **c** is modeled as a one-hot encoding over all cohorts composing a given dataset (for composite datasets 3CA (BC), 3CA (LC)). Our joint distribution with an additional variable **c** decomposes as follows (ignoring log p(**c**)):

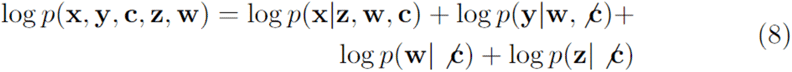

Where we make independence assumptions involving **c** of the form

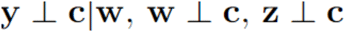

This means that given CNA_d_ latent representations **w**, a cohort variable does not add any further information to CNA profiles **y** (i.e. **w** should contain all necessary information to compute **y**) and that both latent spaces are independent of the cohort. Making the latter assumption allows us to continue using traditional standard normal priors for the latent spaces in this version of the model. The graphical models for ISOMERIC are shown in **S. Fig. 2**.

### Implementation

We implemented ISOMERIC in PyTorch^39^. We use the same architectures and hyperparameters in all experiments. Our architecture for the encoder *p*(**z**,**w**|**x**) takes expression **x** (concatenated with the one-hot encoded **c** in the cohort-encoding model) as input and consists of one hidden 128-dimensional layer, followed by a BatchNorm1d layer with *eps=0*.*001* and *momentum=0*.*01* (following the default SCVI architecture^21^) and a LeakyReLU activation function (chosen due to its suitability for adversarial models). This shared encoder outputs the mean and variance parameters for both **z** and **w**. The expression decoder *p*(**x**|**z**,**w**) takes concatenated samples of **z** and **w** (as well as one-hot-encoded **c** in the cohort-encoding model) as input and mirrors the encoder in its architecture. We model both the adversary *q*(**y**|**z**) and the CNA decoder *p*(**y**|**w**) as a single linear layer to facilitate interpretability and combat overfitting.

We set the latent dimensionality for both **z** and **w** as *D*=10. We trained the models for 200 epochs with Adam optimizer with initial *lr=0*.*01*. We used minibatches of size 256. The hyperparameters were selected in a grid search using validation loss on held out batches (see below). We split the data into patient-aware train and validation sets such that cells from any given patient are either in the training set or validation set. We divided patients in an 80/20% split between the training and validation set for each dataset. When doing this split, we tried to balance the per-patient numbers of cells in the training and validation set to avoid the scenario where all validation patients have a low number of samples. Hyperparameter tuning was performed as per **Supplementary note 1**.

### Robustness to initialization

For each dataset, we trained ISOMERIC over 5 random initializations. ISOMERIC is robust to initialization as it generated consistent cell embeddings in **z** and **w** between random seeds (data not shown). For each dataset, we selected one model out of 5 random seeds to further investigate in this work. We selected each model based on the highest adversarial loss (MSE between the true CNA profile **y** and the predicted **y**_**pred**_ ∼ *q*(**y**|**z**)) on the training set. We focus our selection on the training set here as we are interested in discovering biological insights about our training cohorts, as opposed to achieving high generalization to unseen cohorts.

### scRNA-seq datasets processing

For Peng (PDAC) and Zhou (PDAC) we downloaded Cell Ranger outputs for each tumour sample and read barcode files, feature files, and raw matrices into R. Quality control was performed to remove low quality cells with less than 500 detected genes, 1000 total reads, and mitochondrial gene expression > 20%. Genes with zero detected UMI were filtered out. We used DoubletFinder^40^ o find and remove doublet cells as suggested in documentation and SingleR^41^ to perform cell type annotation for the samples. A multi-cohort PDAC single-cell atlas^42^ was used as the reference for annotation. After SingleR annotation, single-cell labels were manually inspected to collapse “Epithelial acinar cell” and “Epithelial ductal cell” into the “Pancreatic epithelial cell” label.

For all datasets, we additionally filtered genes to protein-coding genes, removed mitochondrial and ribosomal genes, and filtered to genes located on chromosomes 1-22. We filtered cells to tumour cells that had a matched CNA profile (see below). We then retained patients with >100 tumour cells in each dataset. After splitting patients into training and validation sets in each dataset, we removed genes that had no expression in any of the training cells. The final dimensions of all datasets are listed in **Supplementary Table 1**. For Gao et al. (2021) and Pal et al. (2021), cohorts in 3CA (BC), the data was split into multiple files with differing numbers of genes, which we subset to common genes and combined within-cohort. When pooling cohorts into a dataset (3CA (LC) and 3CA (BC)), we again subset to common genes across all cohorts.

When training models with Gaussian likelihood, we applied per-cell library size normalization to 10^4^ cells and log(x+1) transformation. When training models with negative binomial likelihood, we used raw counts.

### Copy number variation estimation with inferCNV

For Peng (PDAC) and Zhou (PDAC) the “normal” reference for inferCNV was constructed by randomly sampling 2000 epithelial cells from normal tissue samples from Peng (PDAC) and two other pancreatic datasets^43,44^ (all normal samples were processed as described in *scRNA-seq datasets processing*). Pancreatic epithelial cells from each tumour sample were combined with the normal epithelial reference independently, such that inferCNV was run on each tumour sample independently. We ran inferCNV with denoising and hidden Markov model subcluster mode activated and with the following parameters: cutoff=0.1, analysis_mode=‘subcluster’, denoise=TRUE, sd_amplifier=3, noise_logistic = TRUE, HMM=TRUE, HMM_type=‘i6’. We used the following inferCNV outputs to classify malignant cells: inferred copy number variation (CNV) in each region and cell groupings clustered by smoothed gene expression. For each sample, we calculated the variance of the regional CNV states in each cell. We then inspected cell clusters for marker gene expression and the CNV variance of cells in them. Clusters with mean CNV state variance > 1 were collapsed into the “Epithelial malignant” compartment and cells assigned to those clusters were labelled as malignant. Clusters with mean CNV state variance <= 1 were collapsed into the “Epithelial normal” compartment.

For SPECTRUM (OV) the authors shared with us the per-patient inferCNV output files. The authors subsampled 500 cells per tumour site (or used all cells if a tumour site had <500 cells) and analyzed those with inferCNV per patient, while sharing a single non-malignant reference across patients. Cells’ metadata contained cell type annotations of multiple cancer cell clusters and two non-malignant ciliated cell clusters, which we used to identify cancer cells.

For 3CA (BC) and 3CA (LC) cohorts, the 3CA maintainers shared with us files containing log-transformed inferred CNAs for 5000 highly expressed genes across cells with >1000 detected genes in each cohort. Cells metadata contained cell type annotations; we used the “Malignant” label to select cancer cells.

### Copy number profile generation

For the three datasets that we had inferCNV outputs for (SPECTRUM (OV), Peng (PDAC), Zhou (PDAC)), we used the output file containing inferred cell subclusters (17_HMM_predHMMi6.leiden.hmm_mode-subclusters.cell_groupings) and the file containing inferred gene copy number (17_HMM_predHMMi6.leiden.hmm_mode-subclusters.pred_cnv_genes.dat). For each patient, we first found an intersection between cells in inferred tumour subclusters and cells identified as malignant (as described above). We then computed chromosome arm-level and band-level CNA profiles (see below) for all tumour cells belonging to a patient. Combining over patients in each dataset, we generated paired CNA profiles for each cell in the dataset.

For arm level CNA profiles, for each subcluster, we extracted inferred gene copy number for genes on each chromosome arm. We mapped hidden Markov models states indicating loss (1, 2) to −1, the state indicating normal copy number (3) to 0, and states indicating gain (4, 5, 6) to 1. If there were no calls for a chromosome arm, its copy number was mapped to 0. We then averaged these re-mapped values within each chromosome arm and recorded a 44-dimensional CNA profile for a given subcluster. All tumour cells assigned to a subcluster were labelled with the subcluster’s CNA profile.

For band level CNA profiles, we annotated each gene that had a CNA call in any subcluster with its chromosome band using ideogram data (retrieved from https://ftp.ncbi.nlm.nih.gov/pub/gdp/) at the resolution of 850 bphs, annotating only those genes that fit within a band in their entirety. We then applied the same mapping of copy number states to −1, 0, 1 indicating loss, neutral, gain as above and averaged these within a band for each subcluster, generating a 811-dimensional CNA profile. If a band didn’t have any calls, its state was mapped to 0. All tumour cells assigned to a subcluster were labelled with the subcluster’s CNA profile.

For the 3CA datasets, as we were provided with log2-transformed CNA estimates, we applied the following transformation to make these values more comparable to our processing of inferCNV outputs for other datasets. We applied a base 2 exponential transformation, standard scaling, and winsorization at [-1,1] to the provided values. We then computed arm- and band-level CNA profiles on the transformed values by averaging them within chromosome arm and band, respectively.

### Baselines

For each dataset, we trained two baselines (PCA and scVI) on the same training sets as ISOMERIC. We trained PCA with 20 components on the training tumour cells and used the learnt projection to embed normal cells. scRNA-seq data was pre-processed with a per-cell library size normalization to 10^4^ cells and log(x+1) transformation. We trained scVI with 20 latent dimensions on the training tumour cells for the maximum of 200 epochs and used the trained encoder to embed normal cells. We used scvi.model.SCVI with n_latent=20 and batch_key set to cohort identity when training on the 3CA (BC) and 3CA (LC) datasets; other parameters were set to defaults. We passed raw scRNA-seq counts to the model’s method setup_anndata() without any pre-processing.

### Benchmarking metrics

To compute the benchmarking metrics, we computed the embeddings of training tumour cells and unseen normal cells (see below) in CNA_i_ and CNA_d_ latent spaces using the best model we selected (across 5 random seeds) for each dataset. We also computed the embeddings for the scVI and PCA baselines. We visualized embeddings with the UMAP projection with hyperparameters set to n_neighbors=20, min_dist=0.3.

Ovarian BAF representation in ISOMERIC: We used SPECTRUM (OV) metadata [] detailing BAF scores for each tumour cell. We clustered tumour cell embeddings in each latent space using sc.tl.leiden with resolution=0.5, n_neighbors=20, min_dist=0.3 and computed ARI between Leiden clusters and discretized BAF score for each annotated chromosome arm. We used Scikit-learn’s^45^ BinsDiscretizer(n_bins**=**5, encode**=**‘ordinal’) to compute the discretized BAF scores. We presented the ARI scores as boxplots across chromosome arms for CNA_i_ and CNA_d_. We tested the difference between the score means in CNA_i_ and CNA_d_ with a T-test without assuming equal variance, using scipy.stats.ttest_ind with equal_var=False and other parameters set to defaults.

Patient/cohort representation in ISOMERIC: To compute ARI with patient or cohort identity, we clustered tumour cell embeddings using sc.tl.leiden with resolution=0.5, n_neighbors=20, min_dist=0.3 and computed ARI between Leiden clusters and the patient/cohort label.

Latent space correlation with CNA: To compute correlation between latent dimensions and CNA profiles, we computed the correlation between every dimension d_i_ ∈ z_1_, …, z_10_, w_1_, …, w_10_ and every chromosome band’s copy number across all embedded tumour cells. We then averaged the absolute values of correlations across chromosome bands for each d_i_, computing the mean absolute correlation with CNA profiles for each latent dimension. We reported these values as boxplots over dimensions in each latent space (CNA_i_, CNA_d_, scVI, PCA).

Normal cell embeddings: We selected normal epithelial cells from all patients included in training and validation sets for Peng (PDAC), 3CA (BC), and 3CA (LC). For Peng (PDAC), we used the labels from our classifier (see *Copy number variation estimation with inferCNV*); for 3CA datasets, we used cell type annotations from the available metadata. As the SPECTRUM (OV) dataset did not include normal epithelial cells, we obtained 91 normal epithelial cells from an atlas of human ovarian ageing^46^ accessed in Human Cell Atlas at https://explore.data.humancellatlas.org/projects/0d4aaaac-02c3-44c4-8ae0-4465f97f83ed. The atlas included snRNA-seq profiling of samples from young and reproductively aged ovaries and included cell type annotations. We used cells from the normal sample “ovary1” as it had the most epithelial cells. We used the same genes that were present in each of our tumour cell datasets, respectively.

We computed the embeddings of normal cells in each latent space using our trained models. Note that as ISOMERIC was never given normal cells in training, all normal cells represent unseen/validation samples. We then computed the variance of normal cell embeddings and tumour cell embeddings in each dimension. We reported these values as boxplots over dimensions for each latent space, separately for tumour and normal cells and across all datasets (outliers were removed to improve clarity).

Quality of CNA decoders: We passed tumour cell embeddings through CNA decoders (adversary *q*(**y**|**z**) and CNA_d_ decoder *p*(**y**|**w**)) and computed the corresponding reconstructed CNA profiles. We then computed the correlation between the per-band copy number in each reconstructed CNA profile and true input CNA profile. We averaged these correlations across bands and reported the mean correlation for CNA_i_ (using reconstructions from adversary *q*(**y**|**z**)) and CNA_d_ (using reconstructions from the decoder *p*(**y**|**w**)) for each dataset.

### Signature and tumour site association analysis for ovarian dataset

For each sample in SPECTRUM (OV), we computed the average embedding in each latent dimension of CNA_i_ and CNA_d_, averaging across all tumour cells in that sample. We then performed one-way analysis of variance (ANOVA) between embeddings in each latent dimension and (i) clinical subtype signature and (ii) tumour site annotation. Both annotations were available from the metadata; tumour site was annotated per sample and signatures were annotated per patient. ANOVA was performed at the sample level (each sample from the same patient was assigned the patient’s signature) and independently for CNA_i_ and CNA_d_ latent spaces. We applied Benjamini-Hochberg correction at 0.05 level across all 20 p-values corresponding to the dimensions of CNA_i_ and CNA_d_. To evaluate association between a selected CNA_d_ dimension (w5) and chromosome bands, we computed correlations between the embedding in that dimension and the copy number profile of each band across all training tumour cells.

### Tumour subtype association analysis for PDAC dataset

Computing basal and classical scores and their association with latent spaces. We used published lists of genes associated with basal and classical PDAC subtypes^35^ to compute the basal and classical score for each training tumour cell in the Zhou (PDAC) dataset. We computed these scores with scanpy’s method sc.tl.score_genes^47^ using counts normalized for library size (to 10^4^), log-transformed, and standard scaled as per recommendations in the documentation. We then averaged over cells in each patient to compute patient-level scores and correlated (Pearson correlation) them with patient-level average embeddings in each latent dimension of CNA_i_ and CNA_d_. We applied Benjamini-Hochberg correction at 0.05 level across all 20 p-values corresponding to the dimensions of CNA_i_ and CNA_d_. We computed the strength of association with basal and classical score for each latent space as the sum of absolute values of correlations across its 10 dimensions.

### Tumour microenvironment association analysis

For each annotated cell type present in our datasets, we computed two metrics: (i) overall proportion: *n*_*t*_ / *N*, where *n*_*t*_ is the number of cells belonging to cell type *t* in a given sample/patient and *N* is the total number of cells in that sample/patient; (ii) immune proportion: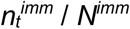, where 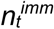 is the number of cells belonging to an *immune* cell type *t* in a given sample/patient and *N*^*imm*^ is the total number of cells across all immune cell types present in that sample/patient.

We computed these metrics for every training sample/patient in each dataset: per sample if a dataset contained multiple samples from the same patient (SPECTRUM (OV) and Zhou (PDAC)) and per patient otherwise. Only samples that had tumour cells were included. For the immune proportion metric, only samples that had immune cells were included. We used author cell type annotations for 3CA (BC), 3CA (LC), and SPECTRUM (OV). For pancreatic datasets, we annotated cell types with SingleR as described in *scRNA-seq datasets processing* and in Yu et al. (2024)^36^. We then correlated (Pearson correlation) these proportions with the average tumour cell embeddings in each latent dimension, aggregated to the sample-or patient-level appropriately. We opted to evaluate this association on a sample level for SPECTRUM (OV) (103 samples) and Zhou (PDAC) (51 samples) to maximize statistical power. This generated 20 correlations (and associated p-values) per cell type and overall proportion metric and 20 correlations per immune cell type and immune proportion metric. We applied Benjamini-Hochberg correction at 0.05 level across all 20 p-values for each cell type and metric independently. Heatmaps summarizing the results only show cell types significantly associated (p_adj_<0.05) with at least one latent dimension and only the significantly associated correlations are shown.

## Acknowledgments

We thank Michael Tyler, Itay Tirosh, Ignacio Vázquez-García, and Sohrab Shah for their help with accessing copy number profiles for the scRNA-seq datasets.

## Contributions

Conceptualization: AS, KRC. Code and analysis: AS, HM, TTJ, CY. Manuscript writing: AS, KRC.

## Data and code availability

All data in this manuscript was previously published elsewhere and is available from public data repositories as outlined above or from correspondence with the original authors. Code presented here for analysis is available at https://github.com/camlab-bioml/2025_isomeric.

