## Supplementary materials for "Learning copy number dependent variation in single tumour cell transcriptomes with deep generative models"

### Supplementary figures

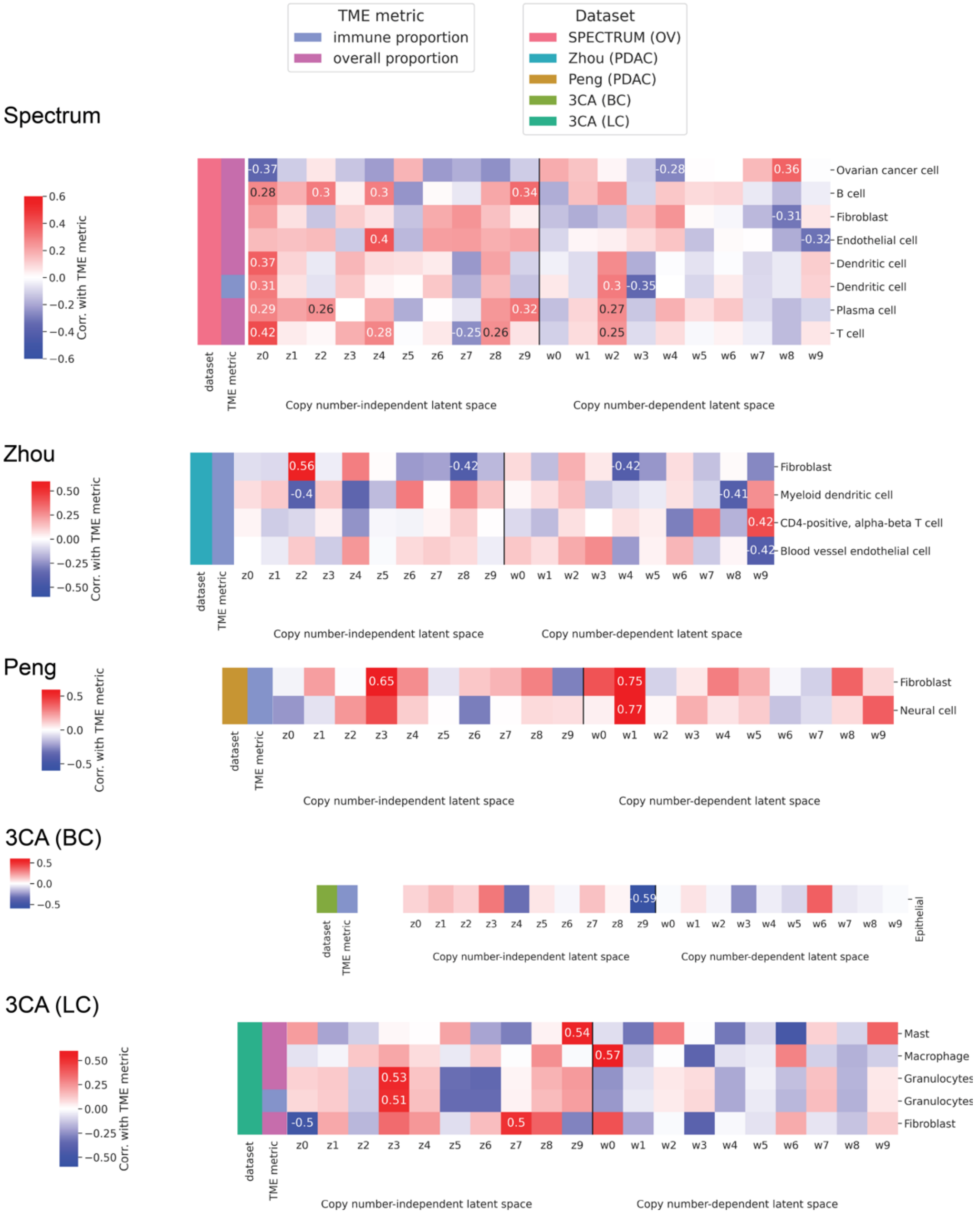

**Supplementary Figure 1**. Association between ISOMERIC latent variables and TME cell type fractions across all datasets.

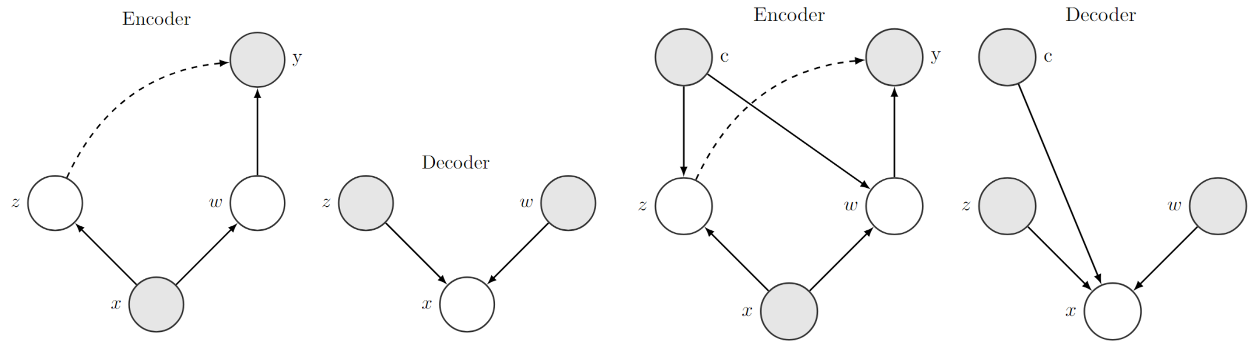


**Supplementary Figure 2**. Left: ISOMERIC graphical model. Right: ISOMERIC graphical model with the cohort encoding variable.

### Supplementary tables

| Dataset | Number of cells | Number of genes | Number of patients |
|---|---|---|---|
| Peng (PDAC) | 23,393 | 16,510 | 23 |
| Zhou (PDAC) | 39,902 | 17,096 | 21 |
| SPECTRUM (OV) | 54,101 | 17,739 | 40 |
| 3CA (BC) | 75,514 | 15,933 | 39 |
| 3CA (LC) | 26,229 | 13,227 | 42 |

**Supplementary Table 1.** The dimensions of generated scRNA-seq tumour cell matrices for the used datasets. The numbers of cells and patients include both training and validation sets.

### Supplementary notes

#### Supplementary note 1: Hyperparameter tuning

We performed hyperparameter tuning in the regime of modeling input data **Y** as chromosome arm-level CNA profiles, with each value indicating the copy number status aggregated over each chromosome arm for a given cell (22 chromosomes * 2 arms = 44 total features). See *Copy number profile generation* for more details. We tuned the following hyperparameters in a grid search for each dataset: (i) number of hidden layers (1, 2) in the encoder and expression decoder; (ii) $M_2$ weight in the minimized loss controlling the adversary (1, 2, 10); (iii) likelihood for gene expression reconstruction (Gaussian, negative binomial), and (iv) the learning rate (0.01, 0.001). Some of the hyperparameter configurations resulted in unstable training regimes with vanishing gradient norms or extremely large loss values; we excluded those runs and those that did not train for 200 epochs from consideration. Results of the tuning experiments are shown in Supplementary Figure 2.

We used the selected hyperparameters (1 hidden layer, Gaussian likelihood, *lr*=0.01, $M_2$=1) for the regime of modeling input data **Y** as chromosome band-level CNA profiles (811 total features). See *Copy number profile generation* for more details. When adapting our training regime to this different number of input features, we modified the loss components computing $M_2$ and $p(\mathbf{y}|\mathbf{w})$ in $M_1$ to have a mean reduction over latent dimensions (as opposed to a sum reduction) and adjusted the weights in the loss as follows. This adjustment kept the overall scale of all loss components similar to the chromosome arm regime.

$$\begin{aligned} m_1(\mathbf{x},\mathbf{y}) = & -\mathbb{E}_{q(\mathbf{z},\mathbf{w}|\mathbf{x},\mathbf{y})}\left[\log p(\mathbf{x}|\mathbf{z},\mathbf{w})\right] - \beta_1 \cdot \left(\mathbb{E}_{q(\mathbf{w}|\mathbf{x})}\left[\log p(\mathbf{y}|\mathbf{w})\right]\right) \\ & + D_{KL}(q(\mathbf{z}|\mathbf{x}) \,||\, p(\mathbf{z})) + D_{KL}(q(\mathbf{w}|\mathbf{x}) \,||\, p(\mathbf{w})) \end{aligned} \tag{9}$$

$$\mathcal{M}_1 = \mathbb{E}_{\mathcal{D}(\mathbf{x},\mathbf{y})}\left[m_1(\mathbf{x},\mathbf{y})\right] \tag{10}$$

$$\begin{aligned} \min \mathcal{M}_1 + \beta_2 \mathcal{M}_2 \\ \max \beta_3 \mathcal{M}_2 \end{aligned} \tag{11}$$

$$\beta_1 = \beta_2 = \beta_3 = 40 \tag{12}$$

For the negative binomial likelihood, we followed the parameterization of Risso et al. (Risso et al. 2018). We modelled gene-specific inverse dispersion parameters as trainable parameters and cell-specific size factors defined as $s_n / 10^4$, where $s_n$ is the library size of cell n. We computed the mean $\mu_{ng}$ for cell n and gene g as the output of the expression decoder multiplied by the size factor of cell n.

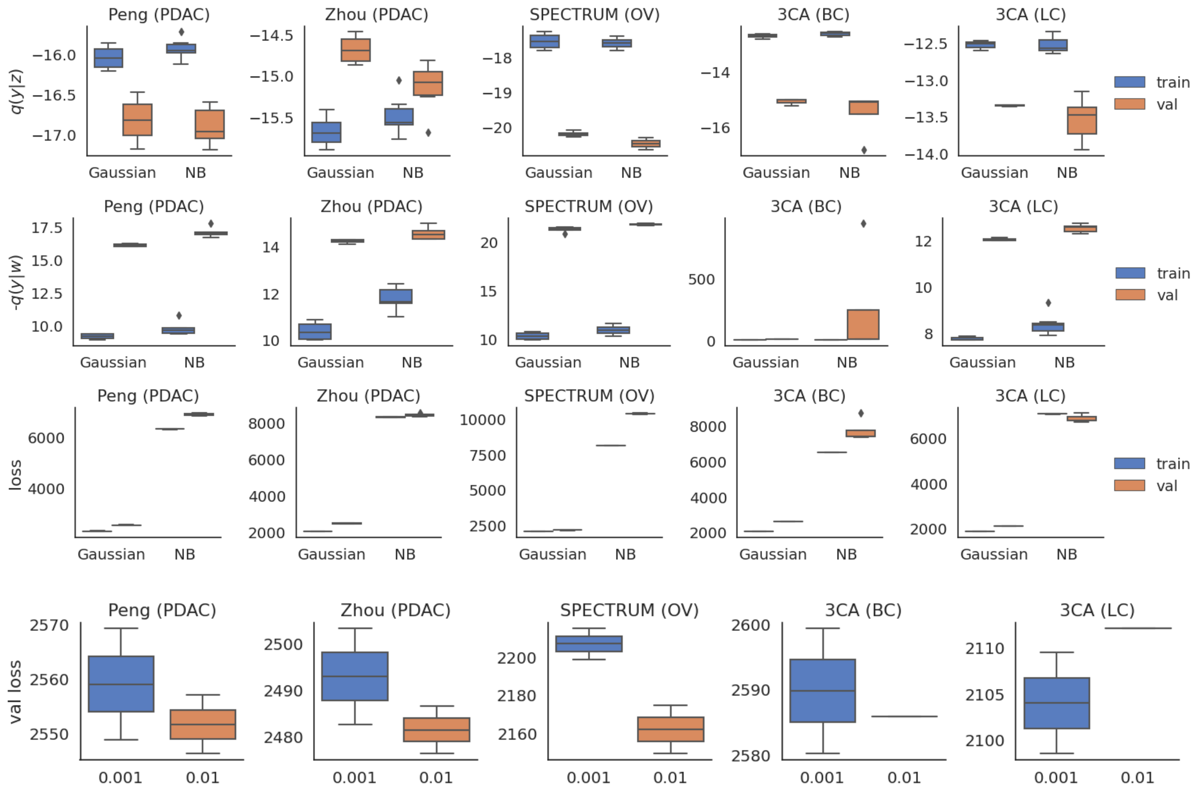


**Supplementary Note 1 Figure 1: Hyperparameter tuning results**. To compare Gaussian and negative binomial likelihood functions, we used *q*(**y**|**z**) and *p*(**y**|**w**) as they compute the conditional likelihood of CNA profiles and are thus comparable across likelihoods. Gaussian likelihood resulted in lower training and validation loss for *p*(**y**|**w**) (top), lower training *q*(**y**|**z**), comparable or lower-variance validation *q*(**y**|**z**) (second from top), and better generalization as quantified by overall loss for all datasets (third from top). For all stable runs with Gaussian likelihood, the number of hidden layers was set to 1 and *lr*=0.01 achieved lower overall validation loss (bottom). Across those parameter settings, $M_2$=1 had a stable run for every dataset and was thus selected.
